# The basolateral nucleus of the amygdala regulates both the predictive and incentive motivational properties of reward cues

**DOI:** 10.1101/755785

**Authors:** Alice Servonnet, Giovanni Hernandez, Cynthia El Hage, Pierre-Paul Rompré, Anne-Noël Samaha

## Abstract

Reward-associated stimuli can acquire both predictive and incentive motivational properties. These conditioned stimuli (CS) can then guide reward-seeking behaviour in adaptive (e.g., locating food) and maladaptive (e.g., binge eating) ways. The basolateral amygdala (BLA) contributes to learning of the predictive value of CS, but less is known about BLA contributions to the incentive motivational properties of appetitive CS. Here we studied the influence of BLA neuron activity on both the predictive and incentive motivational effects of CS. Water-restricted male rats learned to associate a light-tone cue (CS) with water delivery into a port. We assessed the predictive value to the CS by measuring CS-evoked port entries during Pavlovian conditioning. We assessed CS-evoked incentive motivation by measuring lever-pressing for the CS during instrumental responding sessions. During Pavlovian conditioning, we paired CS presentations with photo-stimulation of channelrhodopsin-2 (ChR2)-expressing BLA neurons. This potentiated CS-evoked port entries during conditioning, but suppressed subsequent lever-pressing for the CS. This indicates increased conditioned responding to the CS, but an apparent decrease in incentive motivation for that CS. However, in rats previously naïve to photo-stimulation, pairing BLA-ChR2 stimulations during lever-pressing for the CS intensified responding, indicating enhanced motivation for the CS. Rats did not self-administer BLA-ChR2 stimulations, suggesting that BLA activation does not carry a primary reward signal. Lastly, intra-BLA infusions of d-amphetamine also intensified lever-pressing for the CS. These converging findings suggest that BLA mediated-activity enhances both the predictive and incentive motivational properties of CS, allowing BLA-dependent circuits to guide behaviour in the presence of reward-associated cues.

**SIGNIFICANCE STATEMENT:** Cues paired with rewards can guide animals to valuable resources such as food. Cues can also promote dysfunctional reward-seeking behaviour, as in over-eating. Reward-paired cues influence reward seeking through two major mechanisms. First, reward-paired cues are predictive and thus evoke anticipation of future rewards. Second, reward-paired cues are powerful motivators and they can evoke pursuit in their own right. Here we show that increasing neural activity in the basolateral amygdala enhances both the predictive and motivational effects of reward-paired cues. The basolateral amygdala therefore facilitates cue-induced control over behaviour by both increasing anticipation for impending rewards and making reward cues more attractive.

## INTRODUCTION

Initially neutral cues (sights, sounds or places) that predict rewards such as food and water exert direct control over behaviour. For instance, because reward-paired cues [or conditioned stimuli (CS)] acquire predictive value, they can evoke anticipatory responses allowing animals to prepare for impending rewards. CS can also acquire incentive motivational value (Bolles, 1972; Bindra, 1978), therebying ‘*goading an individual into action*’ (Flagel et al., 2009). In this regard, CS can *i*) elicit approach and attention (Hearst and Jenkins, 1974), *ii*) energize ongoing reward-seeking behaviours (Rescorla and Solomon, 1967), *iii*) trigger the reinstatement of extinguished reward-seeking behaviour (de Wit and Stewart, 1981) and *iv*) reinforce the learning of new instrumental behaviours (Mackintosh, 1974; Cardinal et al., 2002). Via their predictive and motivational properties, CS guide behaviour towards rewards necessary for survival. However, changes in these CS properties can contribute to pathological reward-seeking behaviours (as in addiction) or conversely, low levels of appetitive behaviour (as in depression).

The basolateral amygdala (BLA) regulates both the predictive and motivational effects of CS. BLA lesions (Burns et al., 1993) and optogenetic stimulation of BLA→nucleus accumbens shell neurons (Millan et al., 2017) both attenuate CS-evoked conditioned responses, suggesting disruptions to CS predictive value. The same effect is seen with optogenetic inhibition of either BLA neurons expressing the *Ppp1r1b* gene (Kim et al., 2016) or BLA→nucleus accumbens core neurons (Stuber et al., 2011). The BLA is also necessary for the expression of incentive motivation for CS. Decreasing BLA function with lesions (Cador et al., 1989; Everitt et al., 1991; Brown and Fibiger, 1993; Burns et al., 1993; White and McDonald, 1993; McDonald and Hong, 2004; McDonald et al., 2010), pharmacological agents (Grimm and See, 2000; Kantak et al., 2002; McLaughlin and See, 2003; Rogers et al., 2008; Gabriele and See, 2010) or optogenetic methods (Stefanik and Kalivas, 2013) suppresses behaviour in tasks that measure CS incentive motivation. These studies establish a role for the BLA in appetitive behaviour. However, in these studies, the motivational effects of the CS and UCS were potentially confounded, and/or methods were used (e.g., lesions/pharmacological agents) that do not allow manipulation of neural activity coincident with CS occurrence.

In this context, key questions remain. First, how does increased BLA-mediated neuronal activity during CS presentation influence the predictive and incentive motivational effects of that CS? BLA neurons fire in response to CS presentations during appetitive conditioning (Tye and Janak, 2007; Ambroggi et al., 2008; Tye et al., 2008). The functional significance of this is not fully understood. Second, CS can motivate behaviour through many dissociable psychological processes (Cardinal et al., 2002), what processes might BLA-dependent activity regulate? Increased BLA activity might mediate the specific incentive value attributed to the CS. If so, then increased BLA activity should alter CS motivational properties preferentially when it is explicitly paired with CS presentations. The BLA might also arouse a general motivational state, thereby ‘setting the occasion’ to perform a CS-controlled goal-directed behaviour (Lajoie and Bindra, 1976; Rescorla, 1988). In this case, increased BLA activity should alter incentive motivation for a CS, even when it is explicitly unpaired with CS presentations.

We addressed these questions here using *in vivo* optogenetics combined with Pavlovian and Instrumental conditioning. First, we determined whether photo-stimulation of neurons in the BLA carries a primary reward signal, as assessed by self-stimulation behaviour. We compared self-stimulation of BLA neurons with self-stimulation of adjacent central amygdala (CeA) neurons, as rodents will self-stimulate into the CeA (Seo et al., 2016; Baumgartner et al., 2017; Kim et al., 2017). Second, we determined how photo-stimulation of BLA neurons influences CS predictive value, as assessed by CS-evoked conditioned responses that indicate expectation of the primary reward (Tolman, 1932; Hearst and Jenkins, 1974). Finally, we assessed how photo-stimulation of BLA neurons influences CS incentive motivational effects, by measuring the spontaneous learning of a new instrumental behaviour reinforced by the CS alone (Mackintosh, 1974; Robbins, 1978; Cardinal et al., 2002).

## METHODS

### Animals

Male Sprague-Dawley rats (Charles River Laboratories, Montreal, Canada; 200-275 g on arrival) were housed individually on a 12-h light/dark cycle (lights off at 8:30 a.m.). They were tested during the dark phase of the circadian cycle. Food and water were available *ad libitum*, except in Experiments 3-4, where water access was restricted to 2 h/day. This was to facilitate Pavlovian conditioning using water as the unconditioned stimulus (see below for details). The Université de Montréal approved all procedures involving animals and procedures followed the guidelines of the Canadian Council on Animal Care.

### Intra-cerebral surgery

Rats weighing 325-375 g were anesthetized with isoflurane and placed on a stereotaxic apparatus. For photo-stimulation of amygdala neurons in Exps. 1-3, rats received bilateral infusions of an AAV ChR2-eYFP virus (AAV5-hSyn1-hChR2(H134R)-eYFP; generously provided by Dr. Karl Deisseroth; UNC Vector Core, NC, USA) under the human synapsin promoter into either the BLA (mm relative to Bregma: AP −2.8, ML ± 5.0; mm relative to skull surface: DV −8.4) or CeA (mm relative to Bregma: AP −2.6, ML ± 4.3; mm relative to skull surface: DV −7.9). Control rats received an optically inactive AAV eYFP virus (AAV5-hSyn1-eYFP, UNC Vector Core). Using a glass pipette (tip diameter of ~10 μm) coupled to a Nanoject II (Drummond scientific, PA, USA), we administered 27 microinjections of 36.8 nL each (23 nL/second, at 10-second intervals; total volume of ~1 μL/hemisphere) into each brain region. After the infusions, the glass pipette was left in place for 10 more minutes. In Exp. 4, d-amphetamine was infused specifically into the BLA, or as a neuroanatomical control, into the amygdala without targeting the BLA specifically (referred to as ‘Amygdala’). Guide cannulae (26 GA, model C315G, HRS Scientific, Montreal, Canada) were implanted 2 mm dorsal to the BLA (mm relative to Bregma: AP −2.4, ML ± 5.5; mm relative to skull surface: DV −6.6) or dorsal to the amygdala (mm relative to Bregma: AP −2.3, ML ± 5.1; mm relative to skull surface: DV −6.2). In Exp. 1, the craniectomy was sealed with bonewax (Ethicon, NJ, USA). In Exps. 2-3, an optic fiber implant (~300 μm core diameter, numerical aperture of 0.39; ThorLabs, NJ, USA; glued with epoxy to a ferrule, model F10061F340, Fiber Instrument Sales Inc., Oriskany, NY, USA) was implanted in each hemisphere, 0.2 mm dorsal to the virus injection site. Four to 6 stainless steel screws were then anchored to the skull, and optic fiber implants or cannulae were fixed with dental cement. Optic fiber implants were protected with a sleeve and a dummy. Guide cannulae were sealed with obturators (model C315CD, HRS Scientific). Optogenetic manipulations started at least 4 weeks following virus injection, to allow sufficient viral expression (Zhang et al., 2010).

### *In vivo* electrophysiology

We used *in vivo* electrophysiology to confirm laser-induced action potentials in ChR2-expressing neurons in the BLA and CeA. Anesthetized rats (urethane, 1.2 g/kg, i.p.) were placed inside a Faraday cage on a stereotaxic frame equipped with a body temperature controller. Optrodes were implanted above the BLA and CeA. Optrodes were constructed using an extracellular Parylene coated tungsten electrode (1 MΩ, ~125 μm outer diameter; FHC, Bowdoin, ME, USA) glued with epoxy to an optical fiber (~300 μm core diameter, numerical aperture of 0.39) with a ~0.5 mm offset to ensure illumination of recorded neurons. A reference electrode (insulated silver wire, 0.25 mm diameter) was lowered into the back of the brain close to the cerebellum. The optrode and reference electrodes were fixed with stainless steel screws anchored to the skull and bee wax. The optrode was lowered by hydraulic microdrive into the BLA or the CeA to record single action potentials elicited by laser stimulation (465 nm blue diode laser). Optrodes were linked to the laser via patch-cords built as described in Trujillo-Pisanty et al. (2015).

The signal recorded from each optrode was fed into a high impedance headstage connected to a microelectrode amplifier (Model 1800, A-M Systems, WA, USA). During photo-stimulation, the low- and high-pass filters were set at 300 Hz and 5 kHz, respectively. To reduce the possibility of photoelectric artifacts, we grounded the laser head and patch-cord. Action potentials were displayed on an oscilloscope (Tektronix, Model TDS 1002, OR, USA). The signal was digitalized and stored using DataWave recording (USB 16 channels) and DataWave SciWorks Experimenter Package (DataWave Technologies, CO, USA).

### Pavlovian Conditioning

Training and testing took place in standard operant chambers (Med Associates, VT, USA) where a fan and a house-light were on. Rats had restricted water access for at least 3 days (2 h/day). Starting on the next day, they were trained to associate a light-tone cue [see Fig. 1A; conditioned stimulus (CS)] with water delivery [unconditioned stimulus (UCS); 100 µL] into a recessed receptacle, using Pavlovian conditioning procedures. Levers were always retracted during Pavlovian conditioning sessions. The CS was a light-tone stimulus. It consisted of illumination of two cue lights for 5 seconds, combined with the extinction of the house-light. This was immediately followed by a 1800-Hz, 85-dB tone. The tone lasted 0.18 seconds and was combined with water delivery. The CS-UCS were presented on a variable interval of 60 seconds, 20 or 30 times/session. To measure the predictive effects of the CS, we quantified the number of nose-pokes/minute into the recessed water receptacle when cue lights were on (conditioned stimulus response; CSR) versus the number of nose-pokes/minute during the inter-trial interval (inter-trial interval response; ITIR; see Fig. 1A). As a mathematical index of the predictive value of the CS, we computed a CSR/ITIR ratio for each animal, on each conditioning session.

**Fig 1.**
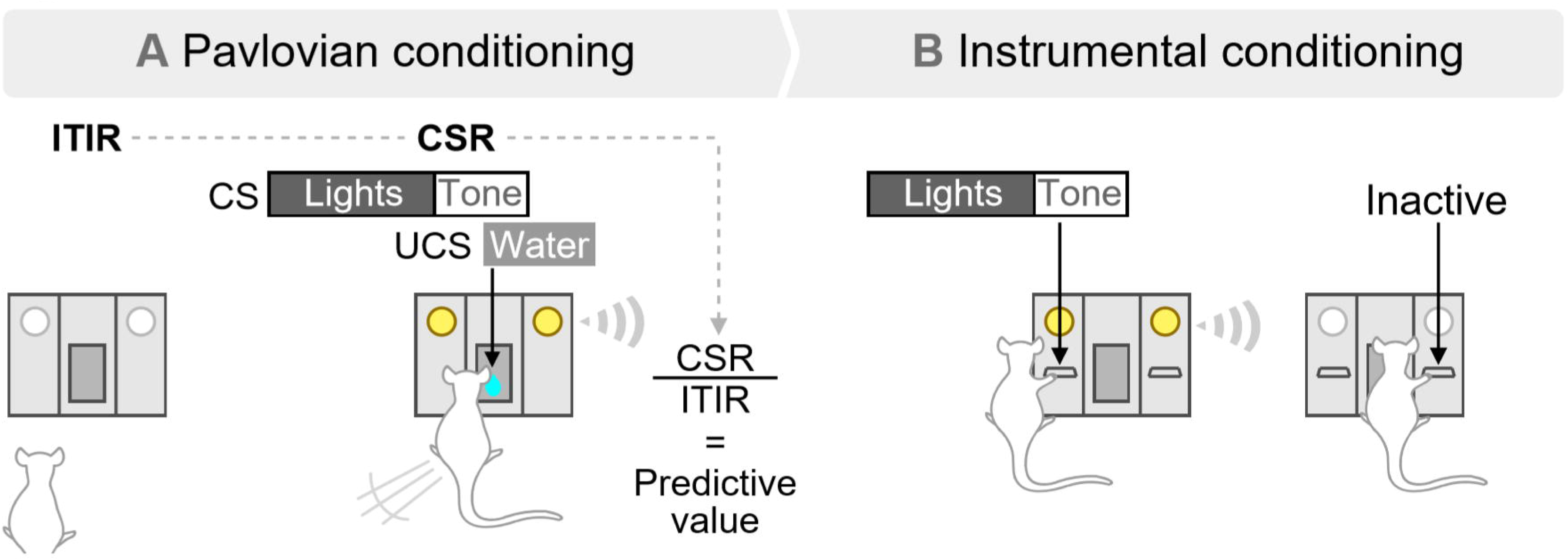
Pavlovian and Instrumental conditioning procedures. (A) During Pavlovian conditioning, rats with limited access to water (2 h/day) learned that a cue (lights + tone, CS) predicts water (100 µl) delivery. The acquired predictive value of the CS was operationally defined as the ratio of the number of nose-pokes/min made during the 5-second cue presentation (conditioned stimulus response; CSR) by the number of nose-pokes/min made during the inter-trial interval (inter-trial interval response; ITIR). (B) After Pavlovian conditioning, rats were given instrumental conditioning sessions during which they were presented with two levers for the first time. Pressing on the active lever produced the CS, while pressing the inactive lever had no programmed outcome.

### Instrumental Conditioning

To assess the incentive motivational effects of the CS, we determined whether after Pavlovian CS-UCS conditioning, rats would spontaneously learn a new instrumental response (lever-pressing) to earn CS presentations, without the UCS. This procedure dissociates incentive motivation for the CS versus for the UCS, because the instrumental response is new and not previously reinforced by the UCS (Mackintosh, 1974; Robbins, 1978; Cardinal et al., 2002). First, rats were placed in the operant chambers for a lever habituation session, during which they could sample the two test levers for the first time. As shown in Fig. 1B, pressing the active lever produced the CS, without water delivery, according to a random-ratio 2 (RR2) schedule. Pressing on the active lever during CS presentation or on the inactive lever had no programmed consequences but was recorded. The lever habituation session ended after 10 active lever presses or 40 minutes. To measure the incentive motivational value of the CS, rats received additional instrumental test sessions. During these sessions, conditions were the same as during the lever habituation session, except that lever presses were not limited. Sessions ended after 20 or 40 minutes. We refer to these sessions as ‘operant responding for the CS’.

### Exp. 1: Effects of photo-stimulation on action potentials in ChR2-expressing BLA and CeA neurons *in vivo*

As shown in Fig. 2A, rats received either the ChR2-eYFP (n = 4) or eYFP (n = 1) virus into the BLA of one hemisphere and into the CeA of the contralateral hemisphere. At least 4 weeks later, rats were anesthetized and *in vivo* neuronal firing was measured following photo-stimulation [squared light pulses of 5 ms delivered at 1, 10, 20 or 40 Hz at 10 mW; based on Huff et al. (2013) and Robinson et al. (2014)]. These are the photo-stimulation parameters used in the behavioural studies below, with frequencies ≤ 20 Hz, at which we observed excellent ChR2 fidelity. Importantly, BLA neurons also fire *in vivo* at frequencies ≤ 20 Hz in behavioural tasks involving reward cues (Tye and Janak, 2007; Ambroggi et al., 2008; Tye et al., 2008).

**Fig 2.**
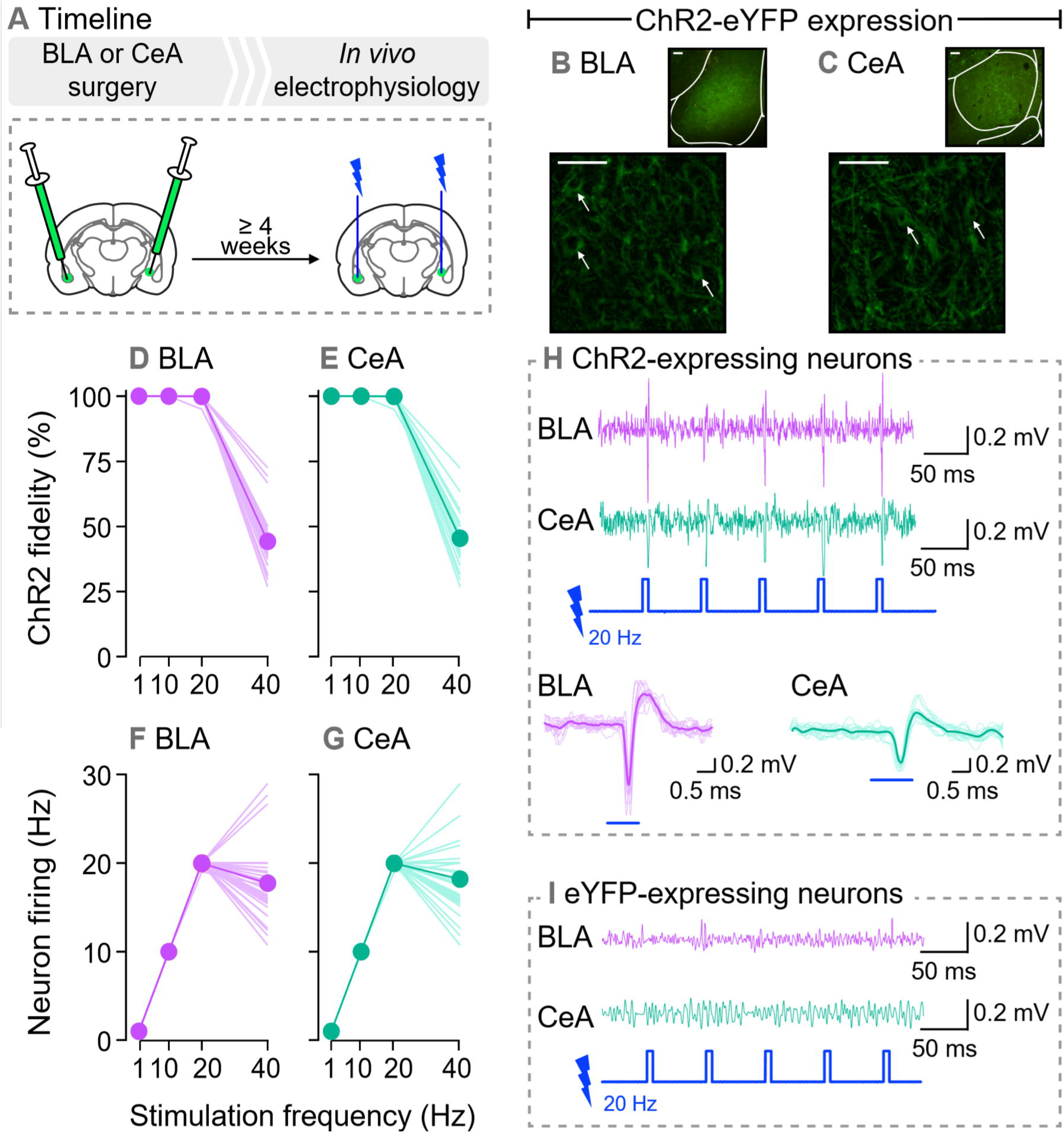
Photo-stimulation reliably induces action potentials only in basolateral (BLA) and central (CeA) amygdala neurons expressing ChR2. (A) In Exp. 1, rats received AAV5-hSyn1-hChR2(H134R)-eYFP (ChR2-eYFP) for transduction and activation of BLA or CeA neurons. Control rats received an optically inactive virus lacking ChR2 (AAV5-hSyn1-eYFP) in the BLA or CeA. At least 4 weeks later, we measured action potentials evoked by photo-stimulation using *in vivo* electrophysiology. (B,C) show ChR2-eYFP expression in the BLA and CeA, respectively (scale bars: 50 µm, arrows indicate cell bodies). When laser-light is delivered, ChR2 reliably induced action potentials in (D) BLA and (E) CeA neurons, with stimulation frequencies ranging between 1-20 Hz. ChR2 fidelity was reduced at 40 Hz. Accordingly, firing frequency of (F) BLA and (G) CeA neurons matches laser stimulation frequency only between 1-20 Hz. Recordings in 4 rats/region; 10 observations/rat. Examples of *in vivo* recordings show that laser-light induces action potentials in (H) ChR2-expressing BLA and CeA neurons but not in (I) eYFP-expressing BLA and CeA neurons.

### Exp. 2: Effects of photo-stimulating ChR2-expressing BLA or CeA neurons on lever-pressing behaviour

If photo-stimulation of BLA or CeA neurons is intrinsically reinforcing, it could reinforce lever pressing behaviour and this would confound interpretation of subsequent results. Thus, here we determined whether otherwise naïve rats would reliably lever press for photo-stimulation of BLA or CeA neurons. As shown in Fig. 3A, rats received bilateral injections of the ChR2-eYFP or eYFP virus into the BLA or CeA. Experimental rats are ChR2-eYFP rats (n = 5/subregion) allowed to lever press for photo-stimulation. Control rats included *i*) rats expressing ChR2-eYFP in the BLA (n = 3) or CeA (n = 2) that could lever press but this did not produce photo-stimulation, and *ii*) rats expressing eYFP in the BLA (n = 3) or CeA (n = 2) and allowed to lever press for photo-stimulations. Throughout the study, lever-pressing behaviour was similar across control groups. Thus, they were pooled together for final analysis (n = 10). Photo-stimulation was bilateral except where noted otherwise.

**Fig 3.**
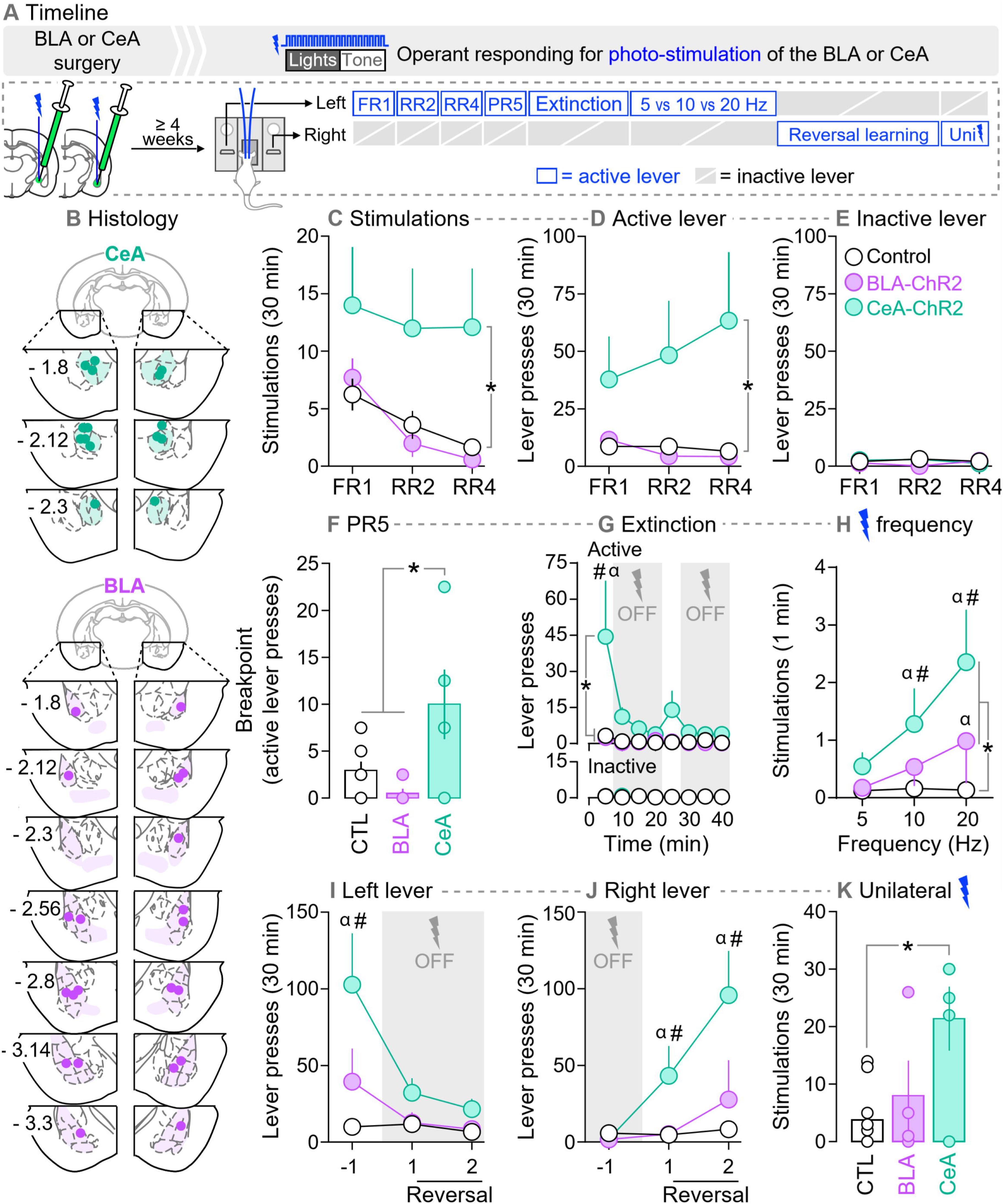
Photo-stimulation of neurons in the central (CeA), but not basolateral (BLA) amygdala is reinforcing. (A) In Exp. 2, rats received AAV5-hSyn1-hChR2(H134R)-eYFP (ChR2-eYFP) or an optically inactive control virus lacking ChR2 (AAV5-hSyn1-eYFP) in the BLA or CeA of both hemispheres. Optic fibers were also implanted bilaterally, above virus injection sites. (B) Estimated optic fiber placements in the CeA and BLA (anteroposterior position is shown in mm relative to Bregma). At least 4 weeks after surgery, rats were allowed to press on two levers. Pressing the active lever produced photo-stimulation of BLA or CeA neurons, paired with presentation of a light-tone cue. Pressing the inactive lever had no programmed consequences. Lever-pressing was measured under different schedules of laser reinforcement and test conditions. (C-D) Fixed ratio 1 (FR1), random ratio 2 (RR2) and random ratio 4 (RR4). (F) Progressive ratio 5 (PR5). (G) Within-session extinction test. (H) Effects of laser stimulation frequency on stimulations earned/min. (I,J) Reversal learning. (K) Effects of unilateral stimulation, under a random ratio 2 schedule of laser reinforcement. \**p* < 0.05. In (G), # *p* < 0.05, versus control rats and BLA-ChR2 rats; α *p* < 0.05, 1^st^ 5-min block versus all other 5-min blocks, in CeA-ChR2 rats. In (H), # *p* < 0.05, versus control rats at the same frequency; α *p* < 0.05, versus 5 Hz within the same group. In (I), # *p* < 0.05, versus control rats in session −1; α *p* < 0.05, versus sessions 1 and 2, in CeA-ChR2 rats. In (J), # *p* < 0.05, versus control rats in the same test session; α *p* < 0.05, versus session −1 in CeA-ChR2 rats. *n*’s = 4-10/group.

As shown in Fig 3A, the rats were allowed to press a lever to obtain a 5.18-second laser stimulation (20 Hz frequency, unless stated otherwise) paired with a 5.18-second presentation of the light-tone stimulus described above. Rats were previously naïve to the light-tone stimulus. During all sessions, active lever presses during photo-stimulation and inactive lever presses had no programmed consequences, but both were recorded. Daily sessions ended after self-administration of 30 stimulations or 30 minutes, unless stated otherwise. First, for at least 2 sessions (1 session/day), pressing the active lever produced photo-stimulation under a fixed-ratio of 1 (FR1) schedule of reinforcement. The rats were then tested under RR2, RR4 and progressive ratio 5 (PR5), with 2 sessions/schedule. Extinction responding was then evaluated during two 40-min sessions, based on Ilango et al. (2014). During minutes 0 to 5 and 20 to 25 of the extinction sessions, lever pressing was reinforced with photo-stimulation under RR2. For the remaining minutes of each session, lever pressing produced the light-tone stimulus, without photo-stimulation. At the 20-min mark, a single, non-contingent photo-stimulation combined with the tone-light cue indicated that photo-stimulation was available once again. Next, we assessed the influence of laser stimulation frequency on lever pressing behaviour during 3 sessions (5, 10 and 20 Hz, one frequency/session/day, counterbalanced). We then assessed reversal learning for 2 sessions during which the active and inactive levers were switched. If photo-stimulation of BLA or CeA neurons is reinforcing, then ChR2-BLA rats and ChR2-CeA rats should stop responding on the newly non-reinforced lever, and increase responding on the newly reinforced lever. Lastly, the rats were given a final test session to determine whether unilateral photo-stimulations are sufficient to reinforce lever-pressing behaviour. The stimulated hemisphere was counterbalanced within each group. After the extinction sessions, one rat in the BLA-ChR2 group was excluded from subsequent testing because of increasing aggressive behavior. In this and subsequent experiments, the experimenter observed each rat during testing. Some rats experienced seizures with repeated photo-stimulation of ChR2-containing BLA neurons (rats in the other groups did not show seizure activity). This is consistent with the amygdala kindling model of epilepsy and neuronal plasticity (Goddard et al., 1969; McNamara et al., 1980; Fisher, 1989). Rats that experienced seizures were eliminated from final data analyses (Exp. 2, n = 0; Exp. 3a, n = 3; Exp. 3b, n = 1) except for one rat in Exp. 3a (see below).

### Exp. 3a: Effects of photo-stimulating ChR2-expressing BLA neurons during Pavlovian CS-UCS conditioning on the predictive and incentive motivational properties of the CS

Exp. 2 showed that rats will reliably lever press for photo-stimulation of neurons in the CeA but not BLA. This indicates that stimulation of CeA, but not BLA neurons is reinforcing. We pursued the following experiments with BLA manipulations only, as the reinforcing effects of CeA photo-stimulation could confound data interpretation. We first determined whether photo-stimulation of BLA neurons during Pavlovian conditioning changes the predictive value of the CS, as measured by the CSR/ITIR ratio described above. As shown in Fig. 4A, a new cohort of rats was prepared for optogenetic manipulations as described above. The rats then received Pavlovian conditioning under one of the following 3 conditions: 1) ‘No Laser’, where the CS was presented alone (ChR2 = 11, eYFP = 5), 2) ‘Paired laser’, where photo-stimulation was paired with each CS presentation (ChR2, n = 3; eYFP, n = 3), and 3) ‘Unpaired laser’, where photo-stimulation and CS presentation were explicitly unpaired, by administering laser stimulation half-way between each CS-UCS presentation (ChR2 = 3). The ‘Unpaired laser’ group served to determine whether increased BLA neuronal activity had to coincide with CS presentation in order to change the predictive effects of the CS. If so, then CSR/ITIR ratios in the ‘Unpaired laser’ group should be similar to those in the ‘ChR2-No Laser’ or eYFP rats. One Unpaired-ChR2 rat had a seizure on session 9. Therefore, the effects of BLA photo-stimulation on CSR/ITIR ratios were analysed on sessions 1-8. This rat was subsequently tested in instrumental conditioning when no photo-stimulations were given (see next paragraph). There were no behavioural differences between ChR2-No laser, eYFP-Paired laser and eYFP-No laser rats under any test condition, and they were pooled into one group (controls, n = 19).

**Fig 4.**
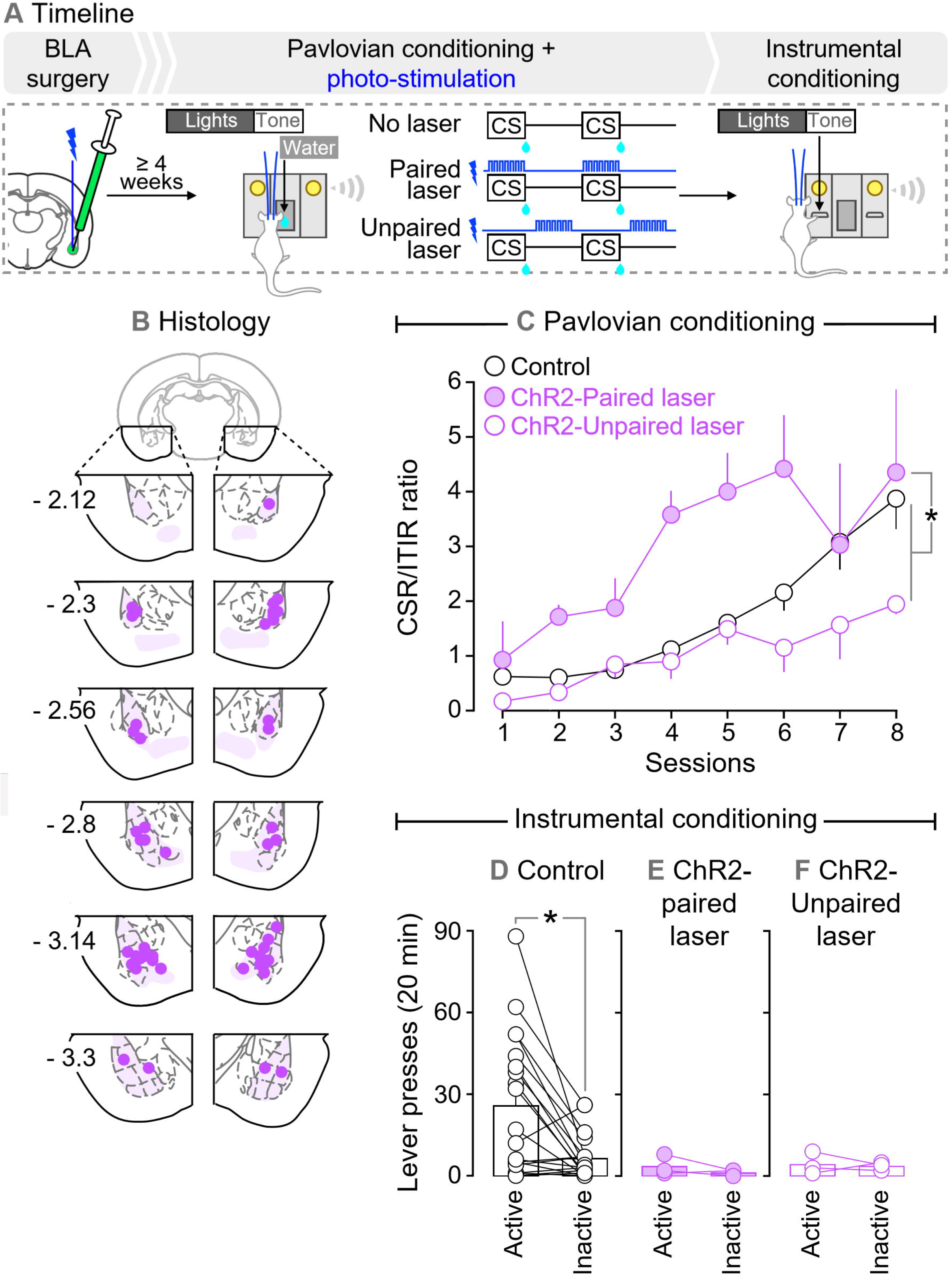
Photo-stimulation of basolateral amygdala (BLA) neurons during Pavlovian conditioning potentiates the predictive value of the conditioned stimulus (CS), but prevents attribution of incentive motivation to that CS. (A) In Exp. 3a, rats received AAV5-hSyn1-hChR2(H134R)-eYFP (ChR2-eYFP) or an optically inactive control virus lacking ChR2 (AAV5-hSyn1-eYFP) in the BLA of both hemispheres. Optic fibers were also implanted bilaterally, above virus injection sites. (B) Estimated optic fiber placements in the BLA (anteroposterior position is shown in mm relative to Bregma). At least 4 weeks after surgery, rats were water-restricted (2 h/day) and received Pavlovian conditioning sessions where a light-tone conditioned stimulus (CS) predicted water (100 µl) delivery (unconditioned stimulus, or UCS). During conditioning sessions, photo-stimulation of BLA neurons was either explicitly paired or unpaired with CS presentation. Control rats (eYFP and ChR2 rats) did not receive photo-stimulations. (B) Paired, but not unpaired photo-stimulation of the BLA enhanced CSR/ITIR ratios (ratio of nose-pokes into the water receptacle during CS presentation versus nose-pokes at other times during the session). This indicates enhanced Pavlovian learning. After Pavlovian sessions, the same rats were given instrumental conditioning sessions, where they could press a lever that produced the CS (active) or a lever that had no programmed outcome (inactive). (D) Rats that had not received photo-stimulations during Pavlovian conditioning pressed more on the active versus inactive lever, indicating incentive motivation for the CS. In contrast, rats that had received BLA photo-stimulation (E) paired or (F) unpaired with CS presentation did not discriminate between the active and inactive levers, suggesting they did not attribute incentive motivation for the CS. *n*’s = 3-19/group. \**p* < 0.05.

We then determined whether pairing photo-stimulation of BLA neurons with CS presentation during Pavlovian conditioning changes subsequent incentive motivation for the CS, as measured by lever pressing for that CS. To evaluate this prediction, ‘Paired laser’, ‘Unpaired laser’ and control rats were allowed to lever press to receive presentations of the CS during a single session.

### Exp. 3b: Effects of photo-stimulating ChR2-expressing BLA neurons during operant responding for a CS

Rats naïve to laser stimulation (control rats from Exp. 3a including eYFP rats and ChR2 rats) received 3 sessions where they could lever press for presentations of the CS paired with BLA photo-stimulation (5, 10 or 20 Hz, one frequency/session, counterbalanced), as shown in Fig. 5A. We then determined whether photo-stimulation of BLA neurons must be explicitly paired with CS presentations to alter operant responding for the CS. If so, then explicitly unpairing photo-stimulation and CS presentation during operant responding for the CS should have no or reduced effects on lever-pressing for that CS. To address this, all rats were given an operant responding session during which photo-stimulation was explicitly unpaired with CS presentation (photo-stimulation applied 3 seconds after each CS presentation).

**Figure 5.**
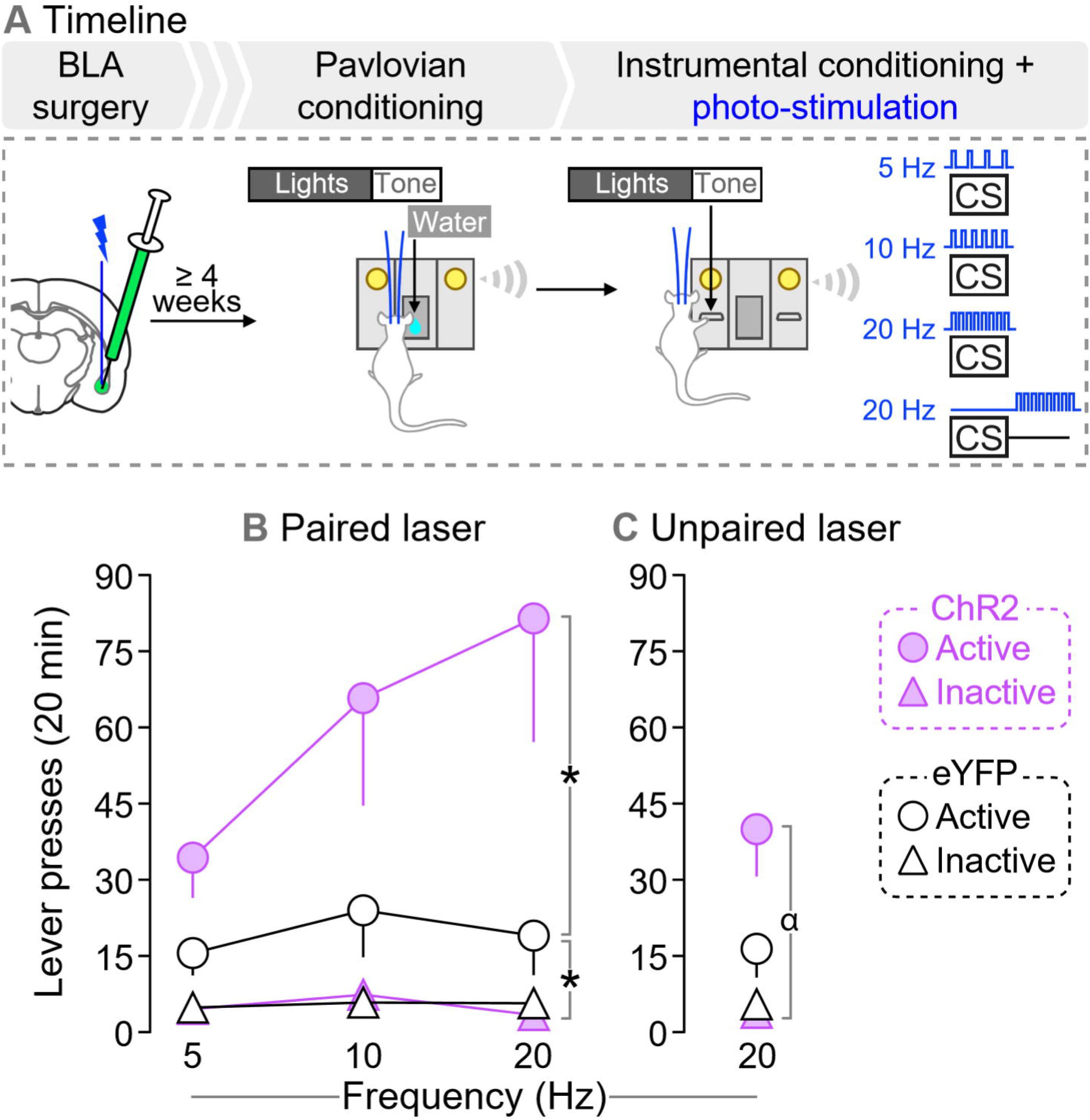
Photo-stimulation of basolateral amygdala (BLA) neurons potentiates the incentive motivational value of a conditioned stimulus (CS). (A) In Exp. 3b, rats that had not received photo-stimulation of BLA neurons during previous Pavlovian CS-UCS conditioning (eYFP rats and ChR2-No laser control rats from Exp. 3a) were used to assess the effects of photo-stimulation of BLA neurons during instrumental responding for the CS. (B) Both ChR2 and eYFP rats pressed more on the active versus inactive lever. ChR2 rats also pressed more on the active lever than eYFP rats did, indicating that photo-stimulation of BLA neurons during CS presentations enhances the incentive motivational value of the CS. (C) When photo-stimulation of BLA neurons is explicitly unpaired with CS presentation, only ChR2 rats pressed more on the active versus inactive lever, but lever-pressing behavior did not differ between ChR2 and eYFP rats. *n*’s =7-8/ group. \**p* < 0.05; α *p* < 0.05, active lever presses versus inactive lever presses of CeA-ChR2 rats.

### Exp. 4: Effects of intra-amygdala d-amphetamine infusions on the incentive motivational effects of a CS

Exp. 3b showed that photo-stimulation of BLA neurons potentiates operant responding for a CS, suggesting that changes in BLA neuron activity influence incentive motivation for CS. Here we sought to extend these findings by using a pharmacological approach to influence BLA neuron activity. Thus, we determined whether injecting d-amphetamine into the BLA also changes operant responding for a CS. We also determined if, within the amygdala, effects of d-amphetamine on incentive motivation for CS are specific to the BLA. To this end, we assessed the effects of infusing d-amphetamine into the amygdala, but without targeting the BLA specifically. We predicted that d-amphetamine infused specifically into the BLA would enhance operant responding for a CS, based on work showing that intra-BLA infusions of d-amphetamine increase cue-induced reinstatement of extinguished cocaine seeking (Ledford et al., 2003). As shown in Fig. 6A, following Pavlovian CS-UCS conditioning, intra-cerebral cannulae were implanted bilaterally. The rats (n = 35) were then given at least 2 weeks to recover. Rats then received a ‘reminder’ Pavlovian conditioning session. Right after this session, rats received intra-cerebral saline infusions to habituate them to the infusion procedure. No behavior was recorded. On the next day, rats received a lever habituation session. Starting on the next day, rats received intra-cerebral saline or d-amphetamine (10 or 30 µg/hemisphere; Sigma-Aldrich, Dorset, UK; 1 injection/day, given every other day) and they were then allowed to lever press for the CS during a 40-min test session. This session length was chosen based on our previous work with intra-nucleus accumbens d-amphetamine injections (El Hage et al., 2015). Each rat received a maximum of 3 intra-cerebral injections to minimize tissue damage, and only 1 d-amphetamine injection. For intracerebral injections, injectors (33 GA, model C315I, HRS Scientific) were inserted to extended 2 mm beyond the cannulae. Microinjections were given in a volume of 0.5 µL/hemisphere and were infused over 1 minute using a microsyringe pump (HARVARD PHD, 2000: HARVARD Apparatus, Saint-Laurent, Canada). Injectors were left in place for an additional minute after the infusion.

**Figure 6.**
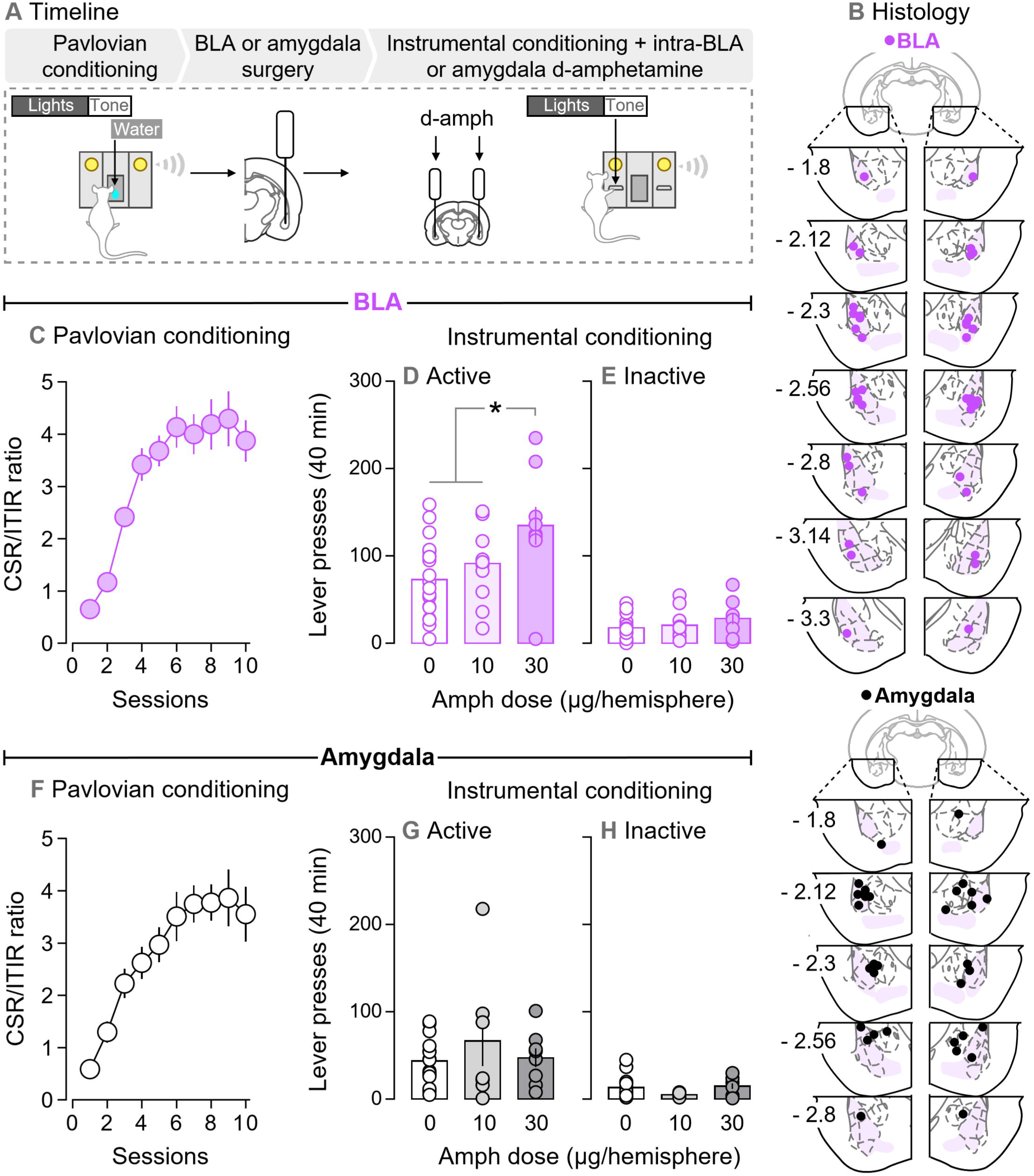
Bilateral infusions of d-amphetamine specifically into the basolateral amygdala (BLA) intensify the incentive motivational value of a conditioned stimulus (CS). (A) In Exp. 4, rats received Pavlovian conditioning. Bilateral cannulae were then implanted specifically into the BLA (‘BLA’ group) or into the amygdala without targeting the BLA specifically (‘Amygdala’ group). (B) Estimated injector tip placements in BLA rats and in Amygdala rats (anteroposterior position is shown in mm relative to Bregma). (C, F) During Pavlovian conditioning, rats reliably learned the CS-unconditioned stimulus contingency, as indicated by higher CSR/ITIR ratios over sessions (ratio of nose-pokes into the water receptacle during CS presentation versus nose-pokes at other times during the session). Next, we assessed the effects of intra-cerebral d-amphetamine infusions (0, 10 or 30 µg/hemisphere) on instrumental responding for the CS. Rats pressed more on (D,G) the active versus (E,H) inactive lever, indicating that the CS acquired incentive motivational value. The highest dose of d-amphetamine (30 µg/hemisphere) significantly potentiated incentive motivation for the CS only if the drug was infused specifically into the BLA. *n*’s = 7-20/group. \**p* < 0.05.

### Histology

In Exps. 2-3, rats were anesthetized with urethane (1.2 g/kg, i.p.) and were transcardially perfused with phosphate buffered saline and 4% paraformaldehyde. Brains were then extracted and kept at room temperature for 1 week in a 30% sucrose/4% paraformaldehyde solution, and then stored at −80 °C. In Exp. 4, rats were anesthetized with isoflurane (5 %), brains were extracted and stored at − 20 °C. Forty µm-thick coronal slices were cut in a cryostat and optic fiber or injector placement was estimated using the Paxinos and Watson atlas (Paxinos and Watson, 1986).

### Statistics

In Exp. 2, mixed-model ANOVA was used to analyse group differences in self-administered photo-stimulations and lever pressing behaviour (Group × Session; ‘Session’ as a within-subjects variable; Group x Time; ‘Time’ as a within-subjects variable; Group x Laser Frequency; ‘Frequency’ as a within-subjects variable). One-way ANOVA was used to analyse group differences in both breakpoint achieved for laser stimulation (PR5 session) and the number of self-administered unilateral stimulations. In Exp. 3a, mixed-model ANOVA was used to analyse group differences in average CSR/ITIR ratios (Group × Session; ‘Session’ as a within-subjects variable). Number of lever presses on the active and inactive levers were analysed using a two-tailed paired *t* test. In Exp. 3b, mixed-model ANOVA was used to analyse group differences in lever pressing for the CS (Group × Session or Lever Type; ‘Session’ and ‘Lever Type’ as within-subjects variables). In Exp. 4, one-way ANOVA was used to analyse CSR/ITIR ratios across sessions. The effects of d-amphetamine on lever pressing were analysed using mixed-model ANOVA (Dose × Lever Type; ‘Lever Type’ as a within-subjects variable). When an interaction and/or main effects were significant (*p* < 0.05), effects were analysed further using Bonferroni’s multiple comparisons’ tests. Values in figures are mean ± SEM.

## RESULTS

### Exp. 1: Effects of photo-stimulation on action potentials in ChR2-expressing BLA and CeA neurons *in vivo*

Figs. 2B-C show ChR2-eYFP expression in the BLA and CeA. As seen in Figs. 2D-E, photo-stimulation of BLA or CeA neurons induced action potentials on average 100 % of the time at 1, 10 and 20 Hz stimulation frequencies. However, at 40 Hz, spike fidelity decreased, and photo-stimulation produced action potentials only ~45 % of the time. In line with these observations, Figs. 2F-G show that the frequency of neuron firing closely matched the frequency of photo-stimulation at laser frequencies ≤ 20 Hz. However, at a stimulation frequency of 40 Hz, BLA and CeA neurons fired at a frequency of only ~18 Hz. This loss of fidelity is in accordance with the kinetic properties of ChR2(H134R), the ChR2 mutant used here. Indeed, when 5-ms pulses are given at a 40-Hz stimulation frequency, pulses are spaced by 20 ms, and this is shorter than the combined opening (~3 ms) and closing (~18 ms) rates of ChR2(H134R) (Lin et al., 2009). Importantly, laser application produced action potentials in ChR2-expressing BLA or CeA neurons (Fig. 2H), but not in eYFP-expressing BLA or CeA neurons (Fig. 2I). Thus, photo-stimulation reliably induced action potentials only in ChR2-expressing BLA or CeA neurons, and spike fidelity was excellent at laser frequencies ≤ 20 Hz. Thus, we used frequencies ≤ 20 Hz in the following studies.

### Exp. 2: Effects of photo-stimulation of BLA or CeA ChR2-expressing neurons on lever-pressing behaviour

Here, we determined whether rats would reliably press on a lever for photo-stimulation of ChR2-expressing BLA or CeA neurons (Fig. 3A). Pressing on the active lever produced photo-stimulation paired with the light-tone cue described above, under FR1, RR2 and RR4 schedules of reinforcement. Pressing on the inactive lever had no programmed consequences.

#### Laser self-stimulation

Fig. 3B shows estimated optic fiber placements in the CeA and BLA. Fig. 3C shows that across different reinforcement schedules, CeA-ChR2 rats self-administered more laser stimulations than control rats (main effect of Group, *F*(2, 17) = 5.5, *p* = 0.014; CeA-ChR2 versus control rats, *F*(1, 13) = 7.5, *p* = 0.017). Accordingly, as seen in Fig. 3D, CeA-ChR2 rats also pressed more on the active lever than controls (main effect of Group, *F*(2, 17) = 5.53, *p* = 0.014; CeA-ChR2 versus control rats, *F*(1, 13) = 7.42, *p* = 0.017). In contrast, BLA-ChR2 and control rats earned a similar number of photo-stimulations and pressed a similar number of times on the active lever (Figs. 3C-D; all *P*’s > 0.05). Presses on the inactive lever did not differ between groups (Fig. 3E; *p* > 0.05), suggesting that photo-stimulation of either BLA or CeA neurons does not have nonspecific motor effects. Fig. 3F shows breakpoints achieved for photo-stimulation under a PR5 schedule of reinforcement. CeA-ChR2 rats reached higher breakpoints relative to BLA-ChR2 or control rats (main effect of Group, *F*(2, 17) = 6.05, *p* = 0.01; CeA-ChR2 > controls, *p* = 0.036; CeA-ChR2 > BLA-ChR2, *p* = 0.013), while BLA-ChR2 rats and controls were not different (*p* > 0.05). Thus, across a range of schedules of reinforcement, rats self-administered cued photo-stimulations of CeA neurons, but not BLA neurons. The findings suggest that photo-stimulation of CeA, but not BLA neurons is reinforcing.

#### Extinction responding

We assessed lever-pressing behaviour under extinction conditions during a 40-min session where photo-stimulation was only available from minutes 0-5 and 20-25, under a RR2 schedule. As shown in Fig. 3G (top panel), BLA-ChR2 rats did not differ from controls during this session (all *P*’s > 0.05). Presses on the inactive lever also did not differ between groups (Fig. 3G, bottom panel; *p* > 0.05). However, Fig. 3G (top panel) also shows that when photo-stimulation was available in the first 5 min of the session, CeA-ChR2 rats pressed more on the active lever relative to controls and BLA-ChR2 rats (Group × Time interaction effect, *F*(14, 119) = 4.14, *p* < 0.0001; main effect of Group, *F*(2, 17) = 7.69, *p* = 0.004; CeA-ChR2 versus control rats, *F*(1, 13) = 10.3, *p* = 0.007; minutes 0-5, CeA-ChR2 > controls, *p* < 0.0001; CeA-ChR2 versus BLA-ChR2, *F*(1, 8) = 5.11, *p* = 0.054; post-hoc comparisons on minutes 0-5, CeA-ChR2 > BLA-ChR2, *p* = 0.0001. No other comparisons were significant). CeA-ChR2 rats also extinguished their lever-pressing behaviour during the extinction session (Fig. 3G, top panel; main effect of Time, *F*(7, 119) = 6.12, *p* < 0.0001; minutes 0-5 vs. each subsequent 5-minute block, all *P*’s < 0.0001). Thus, only CeA-ChR2 rats lever-pressed for photo-stimulation when it was available, and decreased responding when it was not. In contrast, BLA-ChR2 rats and control rats lever-pressed very little, regardless of photo-stimulation availability.

#### Self-stimulation as a function of laser stimulation frequency

Fig. 3H shows the influence of stimulation frequency (5, 10 and 20 Hz) on self-administration of photo-stimulations. Sessions stopped after 30 stimulations or 30 minutes. Most CeA-ChR2 rats took the maximum number of stimulations with higher stimulation frequencies (data not shown). Hence, we analysed the rate of responding (i.e., the number of photo-stimulations earned per minute). Relative to control rats, CeA-ChR2 rats earned more photo-stimulations/minute at 10 and 20 Hz (Fig. 3H; Frequency × Group interaction effect, *F*(4, 32) = 4.08; *p* = 0.009; main effect of Group, *F*(2, 16) = 5.76, *p* = 0.013; CeA-ChR2 rats versus controls, *F*(1, 13) = 10.67, *p* = 0.006; CeA-ChR2 > controls at 10 Hz, *p* = 0.046, at 20 Hz, *p* < 0.0001). CeA-ChR2 rats also earned more photo-stimulations/minute as stimulation frequency was increased (Fig. 3H; main effect of Frequency, *F*(2, 32) = 9.31, *p* = 0.0006; CeA-ChR2 rats, 10 > 5 Hz, *p* = 0.019, 20 > 5 Hz, *p* < 0.0001). BLA-ChR2 rats earned more photo-stimulations/min relative to controls only at the highest frequency tested (main effect of Group, *F*(1, 12) = 6.18, *p* = 0.029; BLA-ChR2 > controls, at 20 Hz, *p* = 0.013). No other comparisons were statistically significant. Thus, compared to control rats, BLA-ChR2 rats earned more photo-stimulations/min at 20 Hz, whereas CeA-ChR2 rats earned more photo-stimulations/min at both 10 and 20 Hz. Furthermore, only CeA-ChR2 rats increased their self-stimulation behaviour with increasing laser frequency.

#### Reversal learning

Here we determined whether photo-stimulation of CeA or BLA neurons supports reversal learning. Fig. 3I shows pressing on a lever that produced the light-tone cue and laser stimulation on session ‘-1’, but not on subsequent sessions. Fig. 3J shows pressing on a lever that did not produce cued laser stimulation on session ‘-1’, but did so on subsequent sessions. As seen in Fig. 3I, CeA-ChR2 but not BLA-ChR2 rats pressed more on the reinforced lever relative to control rats (Group × Session interaction, *F*(4, 32) = 5.46, *p* = 0.002; main effect of Group, *F*(2, 16) = 10.05, *p* = 0.002; CeA-ChR2 versus controls, *F*(1, 13) = 19.47, *p* = 0.0007; CeA-ChR2 > controls on session ‘-1’, *p* < 0.0001), and CeA-ChR2 rats also pressed significantly less on this lever after reversal versus before (main effect of Session, *F*(2, 32) = 11.98, *p =* 0.0001; CeA-ChR2 rats, session −1 > session 1, *p* = 0.0002, session - 1 > session 2, *p* < 0.0001). As seen in Fig. 3J, after lever reversal, CeA-ChR2 but not BLA-ChR2 rats pressed more on the newly reinforced lever relative to controls (Group × Session interaction, *F*(4, 32) = 5.94, *p* = 0.001; main effect of Group, *F*(2, 16) = 8.59, *p* = 0.003; CeA-ChR2 rats > controls, session 1, *p* = 0.033, session 2, *p* < 0.0001), and CeA-ChR2 rats also pressed more on this lever after reversal versus before (main effect of Session, *F*(2, 32) = 11.84, *p* = 0.0001; CeA-ChR2 rats, session −1 > session 1, *p* = 0.035, session −1 > session 2, *p* < 0.0001). In summary, cued photo-stimulation of CeA neurons both reliably reinforced lever-pressing behaviour and supported reversal learning, whereas cued photo-stimulation of BLA neurons did not.

#### Unilateral laser stimulation

Lastly, we determined whether unilateral photo-stimulation of CeA or BLA neurons was reinforcing. Fig. 3K shows that CeA-ChR2 but not BLA-ChR2 rats earned more unilateral laser stimulations relative to controls (main effect of Group, *F*(2, 16) = 6.24, *p* = 0.01; CeA-ChR2 > controls, *p* = 0.009). Therefore, unilateral stimulation of CeA, but not BLA neurons sustains self-stimulation.

In summary, across different schedules of reinforcement, operant testing conditions and photo-stimulation parameters, rats did not reliably self-administer photo-stimulation of BLA neurons. In contrast, rats reliably self-administered photo-stimulation of CeA neurons, indicating that it is reinforcing. These findings show that photo-stimulation of BLA versus CeA neurons has dissociable effects, and that CeA but not BLA neurons carry a primary reward signal.

### Exp. 3a: Effects of photo-stimulating ChR2-expressing BLA neurons during Pavlovian CS-UCS conditioning on the predictive and incentive motivational properties of the CS

#### Pavlovian conditioning

Fig. 4B shows estimated optic fiber placements in the BLA. We first determined the effects of BLA neuron photo-stimulation on the predictive value of a CS (Fig. 4A). Predictive value was measured by analysing the ratio of nose-pokes/minute into the water receptacle when cue lights were on (a conditioned stimulus response, or CSR), versus nose-pokes/minute during the ITI (ITI response, or ITIR). A CSR/ITIR ratio greater than 2 indicates that rats nose-poked twice more during CS presentation than during the ITI. Fig. 4C shows the effects of photo-stimulation of ChR2-expressing BLA neurons on the CSR/ITIR ratio over Pavlovian conditioning sessions. CSR/ITIR ratio progressively increased over sessions in all groups, indicating that rats learned the CS-UCS contingency (Fig. 4C; main effect of Session *F*(7, 154) = 10.19, *p* < 0.0001). Pairing photo-stimulation of BLA neurons with CS presentations (‘ChR2-Paired laser’ group) increased the predictive value of the CS relative to all other conditions (Fig. 4C; main effect of Group, *F*(2, 22) = 3.82, *p* = 0.038; vs. Controls, *F*(1, 20) = 4.84, *p* = 0.04; vs. ChR2-Unpaired laser, *F*(1, 4) = 12.9, *p* = 0.023). No other comparisons were significant. Thus, photo-stimulation of BLA neurons potentiated the predictive value of the CS over time, but only if this stimulation was explicitly paired with CS presentation.

#### Instrumental conditioning

Using the same rats as above, we determined whether having received BLA neuron photo-stimulation during prior CS-UCS conditioning changes the incentive motivational properties of that CS, as measured by the spontaneous acquisition of lever-pressing behavior reinforced solely by the CS. Thus, 2-4 days after the final Pavlovian conditioning session, rats received a single session where they could press a lever that produced the CS alone (no water) or an inactive lever that had no programmed consequences. Fig. 4D shows that only control rats, which *had not received* BLA photo-stimulation during prior Pavlovian conditioning, showed incentive motivation for the CS, as indicated by pressing more on the active versus inactive lever (t(18) = 3.51, *p* = 0.003). To our surprise, rats in the ChR2-Paired laser or ChR2-Unpaired laser groups did not discriminate between the active and inactive levers (Figs. 4E-F; all *P*’s > 0.05). In other words, BLA photo-stimulation during CS-UCS conditioning—regardless of whether the photo-stimulation was paired on unpaired with each CS presentation—inhibits the attribution of incentive motivational value to that CS. Thus, BLA photo-stimulation during CS-UCS conditioning markedly enhanced CS predictive effects (ChR2-Paired laser rats in Fig. 4C), but it prevented attribution of incentive motivation to that CS (Fig. 4E).

### Exp. 3b: Effects of photo-stimulating ChR2-expressing BLA neurons during operant responding for a CS

Exp. 3a showed that photo-stimulation of BLA neurons during CS-UCS conditioning prevents attribution of incentive motivation to that CS. Here, we used rats that had received CS-UCS conditioning without laser stimulation to determine effects of BLA photo-stimulation during subsequent operant responding for the CS (Fig. 5A). This was evaluated in rats with ChR2-expressing BLA neurons (referred to as ‘ChR2’ rats) and rats with eYFP-expressing BLA neurons (referred to as ‘eYFP’ rats). Fig. 5B shows presses on an inactive lever and on an active lever that produced the CS paired with BLA photo-stimulation at different laser frequencies (5, 10 or 20 Hz). Fig. 5B shows that, both ChR2 and eYFP rats pressed more on the active versus inactive lever (main effect of Lever Type, *F*(1, 26) = 18.3, *p* = 0.001; eYFP rats, *F*(1, 12) = 6.31, *p* = 0.027; ChR2 rats, *F*(1, 14) = 12.19, *p* = 0.004). Thus, both groups showed incentive motivation for the CS. In addition, ChR2 rats pressed more on the active lever than eYFP rats (Fig. 5B; main effect of Group, *F*(1, 13) = 5.39, *p* = 0.04. No other comparisons were significant), suggesting that photo-stimulation of BLA neurons potentiates incentive motivation for the CS. Fig. 5C shows lever-pressing behaviour when pressing the active lever produced the CS and photo-stimulation 3 seconds later, such that the CS and photo-stimulation were unpaired. Only ChR2 rats pressed more on the active versus inactive lever (Fig. 5C; Group × Lever Type interaction, *F*(1, 13) = 5.79, *p* = 0.032; main effect of Lever Type, *F*(1, 13) = 20.13, *p* = 0.0006; ChR2 rats, active > inactive lever, *p* = 0.0004). eYFP rats did not discriminate between the levers (*p* > 0.05). No other comparisons were significant. Thus, photo-stimulation of BLA neurons during operant responding for the CS potentiated incentive motivation for that CS, and this effect was stronger when photo-stimulation was explicitly paired with each CS presentation.

### Exp. 4: Effects of intra-amygdala d-amphetamine infusions on the incentive motivational effects of a CS

After CS-UCS Pavlovian conditioning, rats were given instrumental responding tests where they could lever-press for the CS (Fig. 6A). Immediately prior to these tests, rats received bilateral infusions of d-amphetamine (0, 10 or 30 µg/hemisphere) into the BLA or into the amygdala without targeting the BLA specifically. Fig. 6B shows estimated location of injector tips when both cannulae were specifically in the BLA (top) or simply anywhere in the amygdala (bottom). All rats learned the CS-UCS contingency, as indicated by a progressive increase in CSR/ITIR ratio (Fig. 6C; main effect of Session, *F*(9, 171) = 25.35, *p* < 0.0001; Fig. 6F; main effect of Session, *F*(9, 126) = 15.04, *p* < 0.0001). Figs. 6D-E-G-H show that rats pressed more on the active versus inactive lever (Figs. 6D-E; Dose × Lever Type interaction, *F*(2, 37) = 5.31, *p* = 0.009; main effect of Lever Type, *F*(1, 37) = 142.4; *p* < 0.0001; Figs. 6G-H; main effect of Lever Type, *F*(1, 27) = 25.61, *p* < 0.0001). Thus, all rats spontaneously learned a new operant response to produce the CS, indicating that the CS acquired incentive motivational value. D-amphetamine influenced active lever pressing only when infused specifically into the BLA, such that active lever pressing was greatest at 30 µg/hemisphere d-amphetamine (Fig. 6D; main effect of Dose, *F*(2, 37) = 4.5, *p* = 0.018; 30 vs 0 µg, *p* = 0.0002; 30 vs 10 µg, *p* = 0.027). In contrast, d-amphetamine did not alter lever-pressing behaviour in rats that received infusions into the amygdala, without specifically targeting the BLA (Figs. 6G-H; all *P*’s > 0.05). No other comparisons were statistically significant. Thus, intra-BLA d-amphetamine intensified incentive motivation for the CS.

## DISCUSSION

We evaluated the respective contributions of the BLA to the predictive and incentive motivational effects of an appetitive CS. Our data indicate that stimulation of BLA neurons influences both effects. First, we found that photo-stimulation of BLA neurons is not intrinsically reinforcing, whereas photo-stimulation of neurons in the adjacent CeA is. Photo-stimulation of BLA neurons during Pavlovian CS-UCS conditioning enhanced CS predictive value, but only if BLA photo-stimulation and CS presentation were explicitly paired. In contrast, photo-stimulation of BLA neurons during Pavlovian conditioning (whether paired or unpaired with CS presentation) completely abolished the incentive motivational properties normally attributed to the CS, as measured by instrumental responding for the CS. In contrast, if BLA photo-stimulation does not occur during CS-UCS conditioning, but only during later instrumental responding for the CS, this potentiates incentive motivation for that CS. Finally, intra-BLA infusions of d-amphetamine also augmented instrumental responding for the CS, suggesting that a local increase in monoamine neurotransmission is also involved in enhanced conditioned incentive motivation. Thus, altered neuronal activity within the BLA facilitates cue-controlled behaviour by potentiating both the predictive and incentive motivational effects of an appetitive CS.

### Photo-stimulation of CeA, but not BLA neurons is reinforcing

Rats reliably lever pressed for photo-stimulation of CeA, but not BLA neurons, suggesting that CeA neurons carry a primary reward signal. Our findings agree with earlier work showing that electrical stimulation of CeA cells is reinforcing (Prado-Alcala and Wise, 1984; Kane et al., 1991). CeA neurons are mostly GABAergic, but they express different neuropeptides and have different anatomical connections. More recent studies show that stimulation of specific CeA neuron populations can also be reinforcing. These neuronal populations include CeA neurons expressing corticotropin-releasing hormone, somatostatin, neurotensin and/or tachykinin 2 (Baumgartner et al., 2017; Kim et al., 2017), and CeA→medial prefrontal cortex projections (Seo et al., 2016). In contrast, using photo-stimulation of CeA neurons without regards to cell subtype as done here, Berridge and colleagues report that CeA photo-stimulation is not reinforcing (Robinson et al., 2014; Warlow et al., 2017). This could involve the CeA subregion where photo-stimulation was applied. Berridge and colleagues (Robinson et al., 2014; Warlow et al., 2017) implanted optic fibers in the posterior CeA, whereas we implanted in the anterior CeA. Our rats did not reliably self-administer photo-stimulation of BLA neurons (these are principally glutamatergic). Rats will electrically self-stimulate some BLA subregions (Prado-Alcala and Wise, 1984; Kane et al., 1991), and studies using optogenetic methods suggest that self-stimulation depends on the BLA circuit targeted. For instance, photo-stimulation of BLA→nucleus accumbens terminals is reinforcing (Stuber et al., 2011; Britt et al., 2012; Namburi et al., 2015), but photo-stimulation of BLA→medial CeA terminals produces avoidance (Namburi et al., 2015). Via distinct cell types and connections, amygdala nuclei and subregions exert many functions, including both appetitive and defensive behaviours (Gallagher and Chiba, 1996). The hSyn promoter used here confers neuron-specific transgene expression, but it does not target neuron subtypes. Future studies will be important to examine roles of specific CeA and BLA neuron subtypes and projections in appetitive behaviour. As this research unfolds, our results support the idea that while the BLA and CeA are connected and can play similar roles in motivational processes (Wassum et al., 2011), they also have distinct appetitive functions (Corbit and Balleine, 2005; Robinson et al., 2014; Warlow et al., 2017).

### Photo-stimulation of BLA neurons during CS-UCS conditioning enhances the predictive value of the CS but suppresses later incentive motivation for that CS

Pairing photo-stimulation of BLA neurons with CS presentation during Pavlovian conditioning potentiated CS-evoked conditioned behaviour, indicating enhanced anticipation of the UCS. Explicitly unpairing BLA stimulation and CS presentation did not influence CS-evoked conditioned behaviour. Thus, increasing BLA neuron activity when a CS is presented amplifies associative CS-UCS learning. We do not believe this involves changes to representations of the water UCS, because BLA lesions do not alter water consumption (Cador et al., 1989). Instead, enhanced CS predictive value likely involves changes in how BLA neurons represent the CS and/or how they encode the CS-UCS association. This could involve increased activity in BLA→nucleus accumbens projections (Ambroggi et al., 2008; Popescu et al., 2009; Jones et al., 2010b; Jones et al., 2010a; Stuber et al., 2011), BLA→prelimbic cortex projections (Dilgen et al., 2013; McGarry and Carter, 2016; Keefer and Petrovich, 2017) and/or BLA→orbitofrontal cortex projections (McDonald, 1991; Schoenbaum et al., 2003). BLA interneurons could also be involved, as their activity is causal in Pavlovian fear conditioning (Wolff et al., 2014). However, less is known about roles in appetitive conditioning, as used here. Finally, signalling mediated by dopamine, glutamate and/or acetylcholine within the BLA (See et al., 2003; Berglind et al., 2006; Feltenstein and See, 2007) could also be involved in the effects we observed.

Surprisingly, BLA photo-stimulation during CS-UCS conditioning suppressed the attribution of incentive motivation to the CS. After Pavlovian conditioning, control rats (i.e., eYFP rats and ChR2-no laser rats) reliably acquired a new lever-pressing response reinforced by the CS, suggesting that they attributed incentive motivational value to the CS. In contrast, rats that received BLA neuron photo-stimulation during Pavlovian conditioning later did not lever press for the CS, even though they showed *amplified* CS-evoked conditioned behaviour during Pavlovian conditioning. We observed this both when photo-stimulation was explicitly paired or unpaired with the CS. One possibility is that repeated optogenetic stimulation of the BLA is aversive, evoking fear-related emotions (LeDoux, 2000). Should this be the case, our findings suggest that this does not disrupt encoding of the predictive CS-UCS relationship, but it compromises the appetitive motivational state that is necessary for the CS to acquire incentive motivation (Tabbara et al., 2016). BLA neurons also fire in response to CS during appetitive conditioning (Tye and Janak, 2007; Ambroggi et al., 2008; Tye et al., 2008), and it is possible that photo-stimulation of BLA neurons during CS-UCS conditioning disrupts the normal cellular processes involved in assigning incentive motivational value to the CS. Whatever the underlying mechanisms, our findings support the idea that the predictive and incentive motivational properties of CS are dissociable (Flagel et al., 2011; Tabbara et al., 2016), and that BLA neuron activity can play distinct roles in these processes, thereby helping animals decide how to respond to CS in their environments.

### Photo-stimulation of BLA neurons during instrumental responding for a CS enhances incentive motivation for that CS

Our findings suggest that once the CS has been imbued with incentive motivational value through prior association with an appetitive UCS, BLA photo-stimulation amplifies the expression of this incentive motivation (as measured by lever-pressing reinforced by the CS alone). Infusing d-amphetamine into the BLA had the same effect, suggesting that increases in monoamine-mediated neurotransmission in the BLA could be involved (Ledford et al., 2003; Bernardi et al., 2009; Gremel and Cunningham, 2009; Lintas et al., 2011). This extends lesion studies showing that the BLA is necessary for operant responding reinforced by a CS (Cador et al., 1989; Burns et al., 1993). BLA photo-stimulation increased lever-pressing for the CS. This likely does not involve any intrinsically reinforcing effects of BLA photo-stimulation, as our photo-stimulation parameters did not reliably support self-stimulation behaviour. Instead, the BLA stores information about CS value, which is then used to guide behaviour (Cardinal et al., 2002). As such, stimulation of BLA neurons could enhance operant responding for a CS by potentiating the incentive value of the CS itself or of the CS-associated reward representation (Mogenson, 1987; Everitt and Robbins, 1992). At a neuroanatomical level, this could involve recruitment of BLA→prelimbic cortex (Miller and Marshall, 2005; Bishop et al., 2011), BLA→orbitofrontal cortex (Lichtenberg et al., 2017) and/or BLA→nucleus accumbens projections (Everitt et al., 1991; Miller and Marshall, 2005; Stuber et al., 2011; Britt et al., 2012; Namburi et al., 2015).

### Conclusions

Thus, increased neuronal activity in BLA-dependent circuits amplifies control over behaviour by an appetitive cue, and this involves two dissociable psychological mechanisms. A first mechanism involves enhanced CS-UCS associative learning, such that the CS triggers increased anticipation of the impending reward. The second mechanism involves amplified expression of incentive motivation for the CS, such that there is enhanced pursuit of the CS. These findings increase our understanding of how BLA neurons enable adaptive responding to appetitive cues in the environment, and how disruptions in BLA-dependent processes might potentially contribute to disorders defined by too much (e.g., addiction) or too little (e.g., depression) reward pursuit.

## ACKNOWLEDGEMENTS

This work was supported by grants from the National Science and Engineering Research Council of Canada (grant number 355923) and the Canada Foundation for Innovation to ANS (grant number 24326). ANS holds a salary award from the Fonds de la Recherche du Québec-Santé (grant number 28988). We thank Nadia Chaudri, Franca Lacroix, Franz Villaruel, Ivan Trujillo-Pisanty, Jonathan Britt, Sean J. Reed, Mike J.F. Robinson and Kent C. Berridge for generous advice on implementation of *in vivo* optogenetics procedures in our laboratory. We also thank Ellie-Anna Minogianis for technical support.

